# Post-translational regulation of metabolism in fumarate hydratase deficient cancer cells

**DOI:** 10.1101/149716

**Authors:** Emanuel Gonçalves, Marco Sciacovelli, Ana S. H. Costa, Timothy Isaac Johnson, Daniel Machado, Christian Frezza, Julio Saez-Rodriguez

## Abstract

Deregulated signal transduction pathways and energy metabolism are hallmarks of cancer and both play a fundamental role in the process of tumorigenesis. While it is increasingly recognised that signalling and metabolism are highly interconnected, the underpinning mechanisms of their co-regulation are still largely unknown. Here we designed and acquired proteomics, phosphoproteomics, and metabolomics experiments in fumarate hydratase (FH) deficient cells and developed a computational modelling approach to identify putative regulatory phosphorylation-sites of metabolic enzymes. We identified previously reported functionally relevant phosphosites and potentially novel regulatory residues in enzymes of the central carbon metabolism. In particular, we show that pyruvate dehydrogenase (PDHA1) enzymatic activity is inhibited by increased phosphorylation in FH-deficient cells. Our work provides a novel approach to investigate how post-translational modifications of enzymes regulate metabolism and could have important implications for understanding the metabolic transformation of FH-deficient cancers.

## Introduction

Cancer is thought to arise from an abnormal accumulation of somatic mutations in the genome that drive complex and profound alterations of the cellular phenotype (*1*). Among these changes, dysregulated energy metabolism is gaining importance as a hallmark of cancer (*2*). Although some recent work elucidated the genetic underpinning of these metabolic changes (*3*, *4*), whether cancer metabolism is tuned via post-translational changes is still largely unknown. In yeast, several studies have shown that signalling has a broad importance in regulating the activity of metabolic enzymes involved in central carbon metabolism and other peripheral pathways (*5*–*7*). By contrast, regulatory phosphorylation of metabolic enzymes in human cells remains largely uncharacterised.

A particularly well-studied metabolic alteration in cancer is driven by mutations of the metabolic enzyme fumarate hydratase (FH). These mutations cause Hereditary Leiomyomatosis and Renal Cell Cancer (HLRCC) tumours, a cancer syndrome characterised by benign tumours of the skin and uterus, and a very severe and aggressive form of renal cancer (*8*). FH catalyses the conversion of fumarate to malate, a reaction that takes part in the tricarboxylic acid cycle (TCA cycle). FH mutations lead to the impairment of the catalytic activity of the enzyme and thereby to the accumulation of its substrate, fumarate, and to profound metabolic changes that we and others have extensively characterised (*8*–*10*). Yet, whether these metabolic changes are interconnected with upstream signalling processes has not been investigated.

Here, we performed an integrative analysis to investigate at a genome-scale level the regulatory interactions between metabolism and signalling using cell lines derived from an HLRCC tumor, UOK262, and the FH reconstituted counterpart, UOK262pFH, which we previously generated (*11*). In particular, we characterised signalling and metabolic changes in HLRCC cell lines by designing and acquiring phosphoproteomics, proteomics and metabolomics measurements (Figure 1). These data-sets allowed us to study the molecular adaptations of signalling and metabolism driven by the loss-of-function of FH in HLRCC using a computational framework that integrates phosphoproteomics with *in silico* estimated metabolic flux rates. Pairing the metabolomics modelling with phosphoproteomics measurements allowed us to identify putative-regulatory phosphorylation-sites in metabolic enzymes involved, mostly, in the central carbon metabolism. Notably, we experimentally validated that phosphorylation of pyruvate dehydrogenase E1 component subunit alpha (PDHA1) regulated metabolism in FH-deficient cells. In summary, we present a novel computational and experimental approach to systematically identify putative regulatory phosphorylation-sites in metabolic enzymes. This approach could reveal novel regulatory networks in the metabolic transformation of cancer, with important implications for cancer therapy.

**Figure 1.**
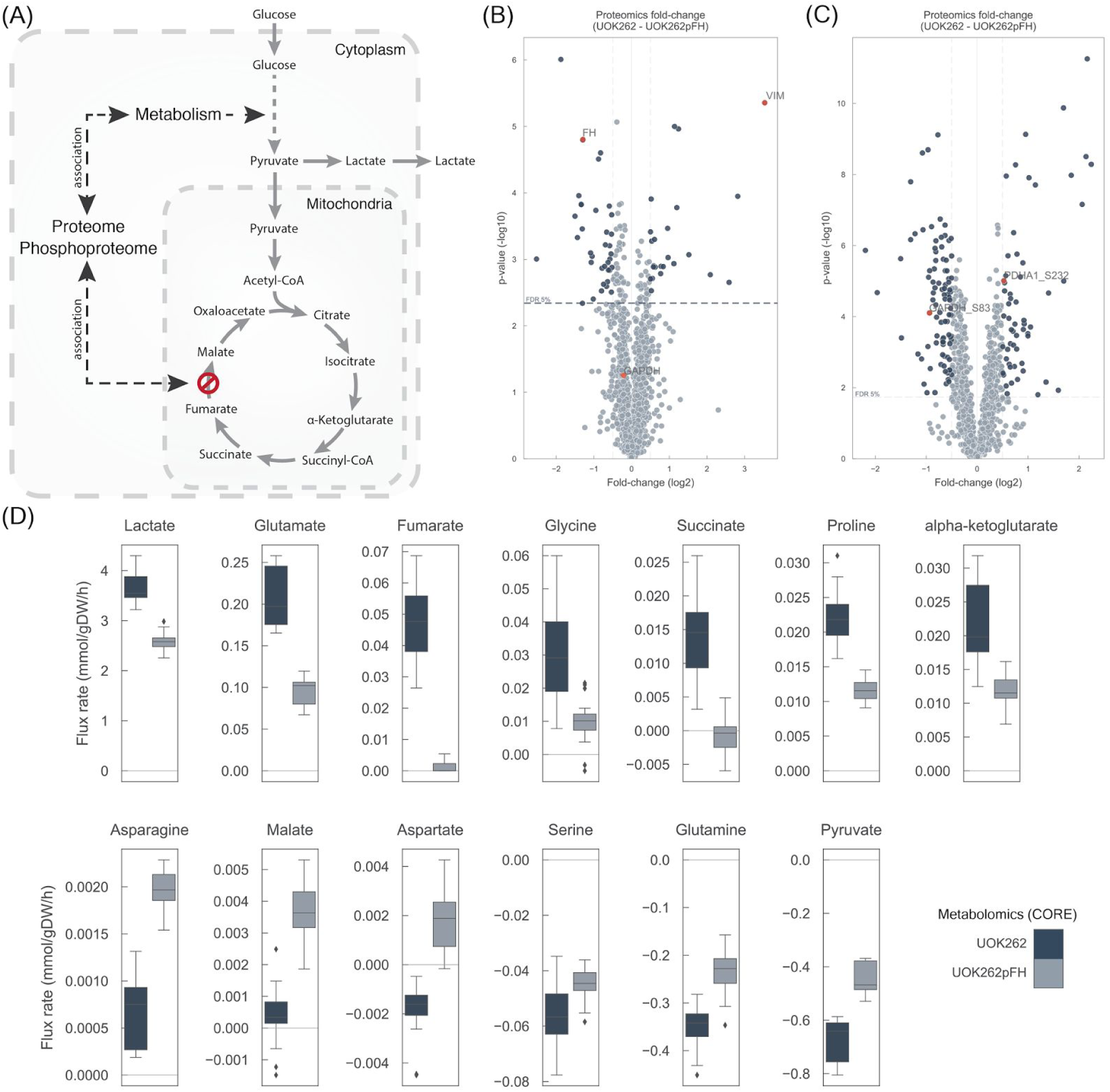
Molecular characterisation of HLRCC derived UOK262 and UOK262pFH cell lines. A) Diagram depicting the potential molecular implication of fumarate hydratase deletion in the proteome and phosphoproteome and subsequent regulatory implications in metabolism. B) Proteomics differential analysis between UOK262 and UOK262pFH. C) Differential phosphoproteomics analysis of phosphosites. D) Consumption and release (CORE) metabolomics experiments quantifying exchange rates (mmol/gDW/h). All the metabolite rates shown are significantly different (FDR < 5%) between UOK262 and UOK262pFH cells.

## Results

### Characterisation of the (phospho)-proteome of human FH-deficient cells

We started our investigation by characterising the proteome and the phosphoproteome of human FH-deficient UOK262 cells and their FH-complemented counterpart, which we previously generated (*10*, *11*). Proteomics experiment covered a total of 1,468 unique proteins (Supplementary Table 1) and, in agreement with FH loss, FH was underexpressed in UOK262 cell lines (Figure 1B). Consistent with previous results, vimentin (VIM) is a mesenchymal marker and was identified as a top-expressed protein (*10*). Reproducibility of the measurements was assessed with unsupervised hierarchical clustering where replicates showed higher correlation coefficients than all the pairwise comparisons (Supplementary Figure 1). Proteomics showed agreement with the RNA-seq transcriptomics measurements available for the same cell lines (spearman’s rho (r) of 0.43, *p-value* = 1.7e-63) (*10*) (Supplementary Figure 2A). Some proteins displayed a disagreement between the protein abundance and the transcript expression, reflecting different types of regulatory mechanisms occurring at post-transcriptional and post-translational levels, consistent with previous reports (*11*).

To study post-translational modifications by phosphorylation we characterised the phosphoproteome of these cell lines. In total, we measured 1,360 unique single phosphorylated phosphosites, mapping to 812 unique proteins (Figure 1C) (Supplementary Table 1). Similarly, to the proteomics measurements, VIM also showed a significant increase in phosphorylation in the UOK262 cell lines, although these changes are associated with the increase in protein abundance. Metabolic enzymes displayed significant changes between UOK262 and UOK262pFH, in particular PDHA1 and GAPDH (Figure 1C). Specifically, 56% (23/41) of the phosphosites in metabolic enzymes display significant changes (FDR < 5%), and these map to 20 unique enzymes. This supports the idea that metabolic enzymes are regulated by phosphorylation in FH-deficient UOK262 cell lines.

### Genome-scale metabolic modelling

To investigate how phosphorylation of metabolic enzymes could regulate metabolism, we computed the intracellular metabolic fluxes of FH-deficient cells using genome-scale reconstruction of human metabolism (*12*–*14*). We constrained the human genome-scale model Recon 2.2 (*14*) with consumption/release (CORE) measurements (Figure 1D), growth rates of UOK262 and UOK262pFH (Supplementary Table 1) and the FH loss status (Figure 2A).

**Figure 2.**
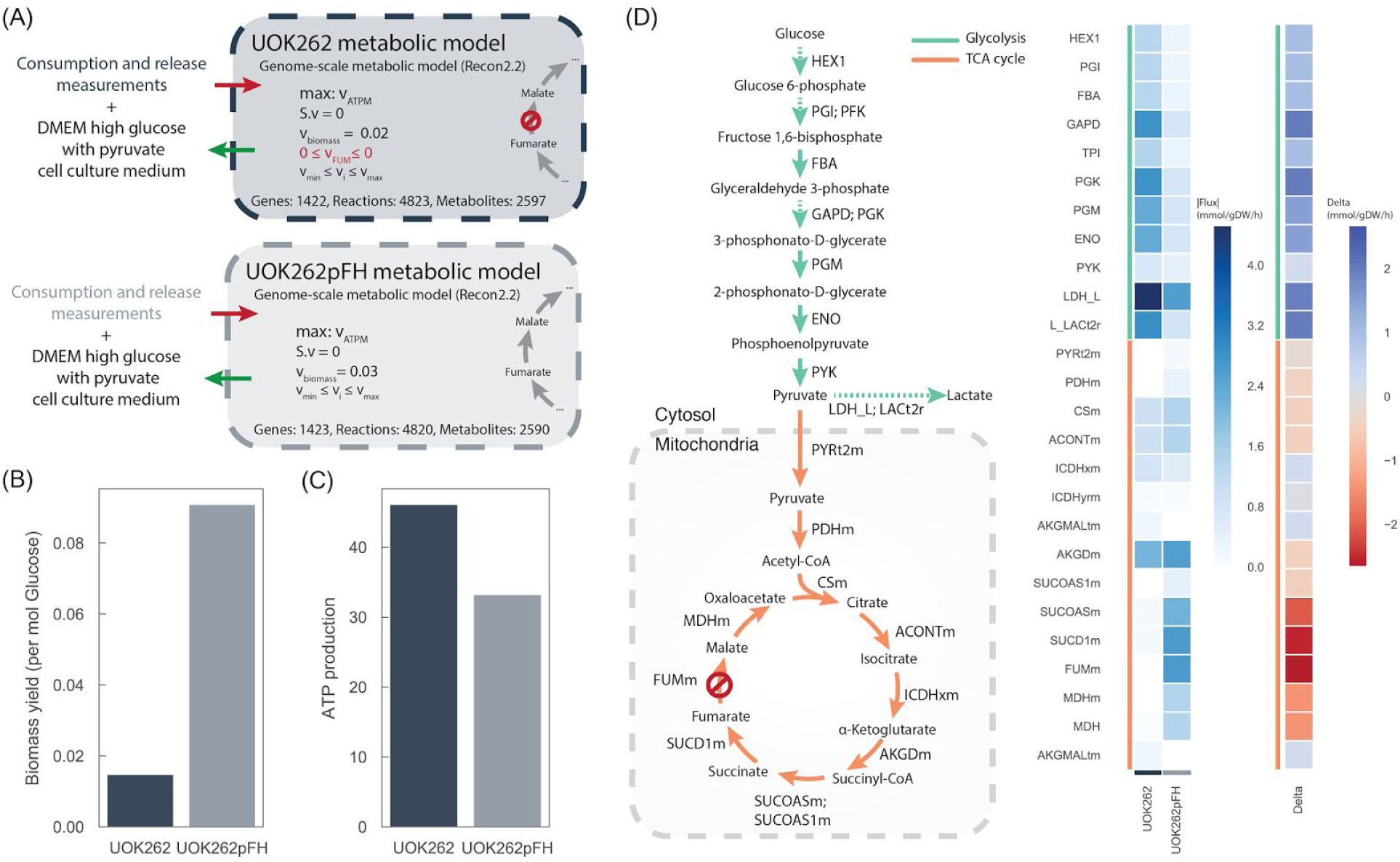
Genome-scale metabolic modelling of UOK262 cell lines. A) Diagram depicting the different constraints used in Recon 2.2 to obtain the condition-specific, UOK262 and UOK262pFH, metabolic models. B) Biomass yield per mol of glucose intake calculated from experimental measurements. C) Maximum ATP production of both models. D) Flux distributions of glycolysis and TCA cycle pathways reactions estimated by maximising ATP production using pFBA.

We used liquid chromatography coupled to mass spectrometry (LCMS)-based to measure CORE rates of 25 metabolites involved in central carbon metabolism, out of which 13 showed significant changes (Figure 1D) (Supplementary Table 1). An unsupervised hierarchical clustering showed that UOK262 and UOK262pFH clustered separately (Supplementary Figure 1). UOK262 cells displayed increased lactate secretion (Figure 1D), and while not significant at an FDR 5% (FDR < 10%) they also showed increase of glucose consumption (Supplementary Table 1), in line with aerobic glycolysis in FH-deficient cells (*11*).

A recent version of the general human genome-scale metabolic model (*14*) was used to generate specific models for UOK262 and UOK262pFH cell lines separately. FH loss in UOK262 cells was represented by limiting the flux rate of its catalysed reactions to zero, while in the UOK262pFH cells they remained unaltered. The composition of the cell medium was also used to restrict the metabolites available for consumption (see Methods). The metabolic models were then constrained using the CORE rates, hence generating two context-specific metabolic models (Figure 2A). Parsimonious FBA (pFBA) (*15*) was then used to simulate the two metabolic models using ATP production as the objective function (Supplementary Table 2). UOK262 cells showed decreased biomass yield, suggesting that the impairment of the mitochondrial function by FH deletion and the increased levels of glucose intake does not lead to augmented growth rate (Figure 2B). The models predicted increased levels of energy production of UOK262 cells (Figure 2C). Of note, the total amount of ATP production in UOK262 was greater than what would be expected by glucose uptake alone, suggesting that other carbon sources are utilised by these cells. This is supported by the measured increased uptake of glutamine.

Glycolytic reactions displayed increased fluxes and increased levels of lactate secretion in UOK 262 cells (Figure 2D), in line with previous observations (*11*). Interestingly, the models also captured the impaired mitochondrial activity of UOK262 cells. Pyruvate dehydrogenase (PDHm) reaction shows decreased intake of pyruvate into the mitochondria and FH inactivation leads to decreased metabolic activity of several reactions in the TCA cycle, e.g. MDHm, SUCD1m. While, these models do not account for intracellular accumulation and depletion of metabolites concentration these results are in line with the accumulation of fumarate, succinate, and succinyl-coa as aKGDm displays consistent activity. Of note, increase of fumarate is a clear biochemical feature of FH-deficient cells (*11*). In summary, these models recapitulate several known metabolic phenotypes of these cell lines and also offer the possibility to explore at a genome-scale level the metabolic adaptations of UOK262 to the reactivation of FH.

### Post-translational and post-transcriptional regulation of metabolism

Having evaluated the predictive capacity of the metabolic models, we then used the *in silico* metabolic fluxes together with the proteomics and phosphoproteomics data-sets to explore potential regulatory mechanisms of metabolism. An exploratory enrichment analysis (*16*) of the proteomics revealed several Gene Ontology (GO) terms (*17*) of different biological processes to be significantly enriched (Figure 3A) (Supplementary Table 2). In particular, processes involving cellular filament and cytoskeleton were identified to be significantly up-regulated in UOK262. This result is consistent with the increased motility of UOK262 cells, which has also been associated with epithelial to mesenchymal transition (EMT) (*10*). Several GO terms related with mitochondrial processes, such as respiratory chain complexes, were down-regulated, consistently with the metabolic models predictions and previously observed decreased mitochondrial activity (*10*). We then assessed if protein abundance changes in metabolic enzymes could be related with metabolic flux changes predicted by the model (Supplementary Figure 2B). Correlation analysis showed no significant relationship (Spearman’s r = 0.12, p-value = 5.21e-01), suggesting that enzyme abundance is insufficient to determine metabolic fluxes, which is consistent with the limited success of previous approaches to interpret metabolism using transcriptomics and proteomics data (*18*). Nevertheless, several metabolic pathways displayed a consistent profile at protein and flux level. For example, the TCA cycle has decreased protein abundance and metabolic flux in UOK262. Intriguingly, glutamate metabolism shows an increase in the abundance of metabolic enzymes and decrease in metabolic flux of the whole pathway in UOK262 (Supplementary Figure 2B), and this was not in agreement with the measured increase in glutamate secretion and glutamine intake (Figure 1D). We validated this unexpected finding performing a separate ^13^C-glutamine labeling experiment and found that indeed, whilst these cells do not accumulate glutamate (*19*), they release glutamate in a time-dependent fashion, and that glutamate is predominantly generated by glutamine (See Supplementary Figure 2C). These observations suggest a broad regulatory role of post-transcriptional changes in the central carbon metabolism of UOK262 cells.

**Figure 3.**
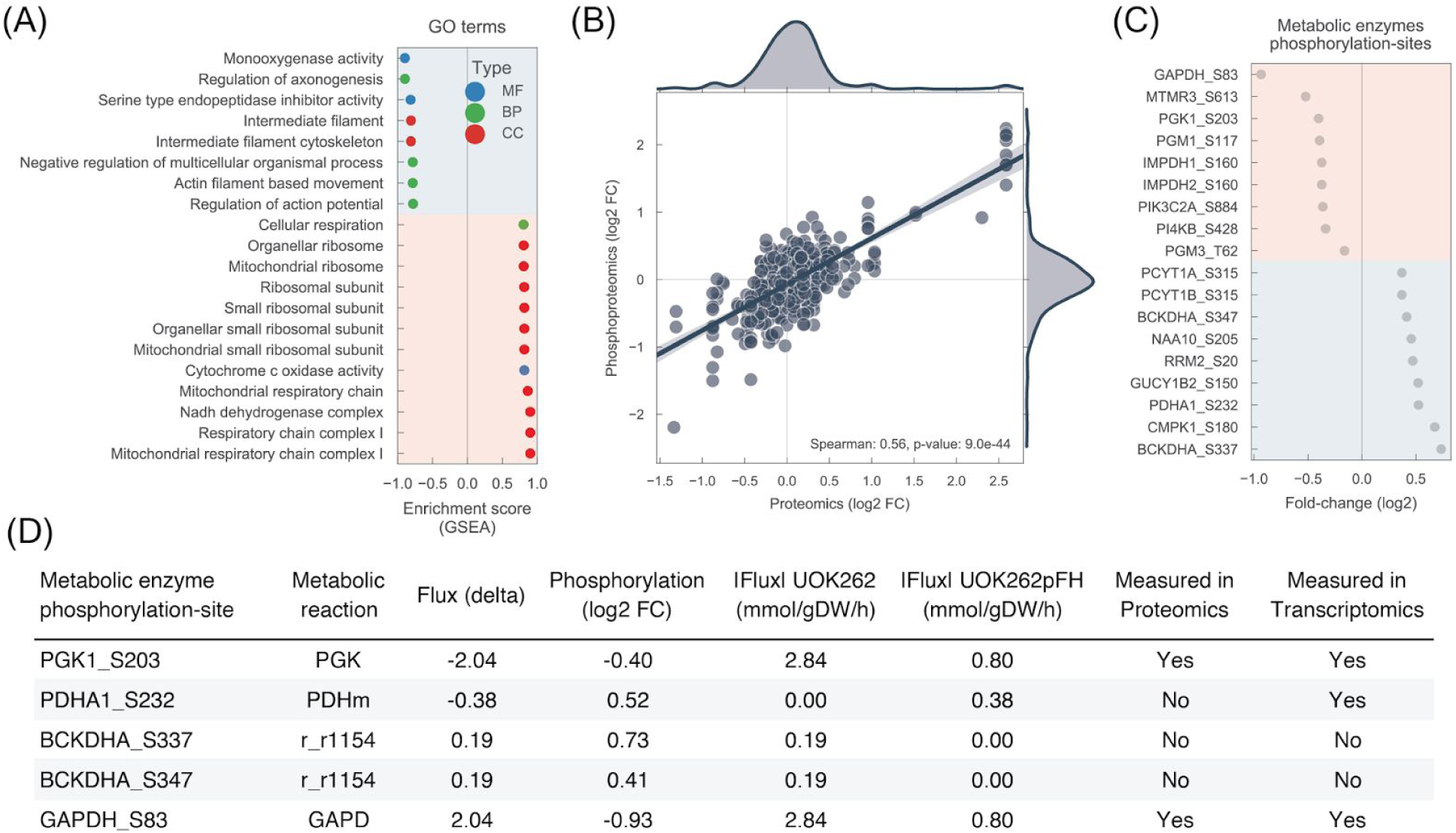
Post-translational regulation of metabolism in FH-deficient cells. A) Top significantly enriched GO terms found in the proteomics data-set (UOK262 - UOK262pFH). Red background denote GO terms that are down-regulated and blue background denotes up-regulated. MF: molecular function; CC: cellular component; BP: biological process. B) Correlation between proteomics and phosphoproteomics measurements. C) Phosphorylation-sites located in metabolic enzymes for which the protein abundance is either not changing significantly or it was not measured. D) List of putative regulatory phosphorylation-sites in metabolic enzymes. Candidates were selected from C) and sorted according to the absolute metabolic flux change (flux delta). The top 5 are shown.

Next, we set to find potential regulatory phosphorylation sites of metabolic enzymes. We first assessed how related are the phosphorylation measurements to the respective abundance of the protein. As expected, phosphosites fold-changes were tightly correlated with the protein abundance (spearman’s r = 0.56, p-value = 9.0e-44) (Figure 3B). To focus only on phosphorylation changes that are independent of the protein abundance we only considered residues that change significantly in phosphorylation but not in abundance. This process allowed us to obtain a list of 18 phosphorylation-sites in metabolic enzymes that show significant changes in phosphorylation in FH-deficient cells (Figure 3C). Consequently, we enquired which of these phosphosites are more likely to have a regulatory role in the metabolic enzymes using the *in silico* metabolic fluxes measurements as a proxy for enzymatic activity. Using the genomic annotation in the metabolic model we mapped the selected phosphosites in the metabolic enzymes to the reactions they catalyse, covering 20 phosphosite-reaction interactions (Supplementary Table 4) (Figure 3D). With our analysis we recapitulated 3 previously reported regulatory residues in PhosphositePlus (*20*), PDHA1_S232, PGK1_S203 and RRM2_S20. Among the top changing reactions with matched phosphorylation changes is PDHA1, where reaction PDHm shows decreased metabolic flux and increased phosphorylation in S232. This result is consistent with current literature that shows that increased phosphorylation in any of the serine residues in positions 232, 293 and 300 of PDHA1 inactivates the enzyme, and its function is only restored when the residues have been dephosphorylated (*21*–*23*). We further assessed the phosphorylation changes in UOK262 and UOK262pFH cells of the other reported regulatory residue, S293, by Western Blotting (WB). Similarly to S232, S293 phosphorylation is decreased in UOK262pFH compared to UOK262 cells (Figure 4A). Moreover, as PDHA1 was not detected in the proteomics data-set, we measured its abundance also with WB, and confirmed that there are no significant changes in its abundance between UOK262 and UOK262pFH (Figure 4B). These results support the hypothesis that PDHA1 activity is regulated by post-translational modifications. We then validated the predicted inactivation of PDHm by measuring the conversion of glucose-derived pyruvate to citrate, a two-step reaction that involves PDH-mediated conversion of pyruvate to acetyl-CoA and the condensation of acetyl-CoA with oxaloacetate to generate citrate. To this aim, cells were incubated with ^13^C_6_-glucose and the isotopologue distribution of pyruvate and citrate was analysed by LC-MS (schematic in Figure 4C). Whilst glucose uptake is increased in FH-deficient cells (*24*) (Supplementary Figure 2D), the incorporation of glucose-derived molecules into citrate is significantly reduced in FH-deficient cells, consistent with the inhibition of PDH activity and the prediction of the model based on phosphoproteomics data (Figure 3C).

**Figure 4.**
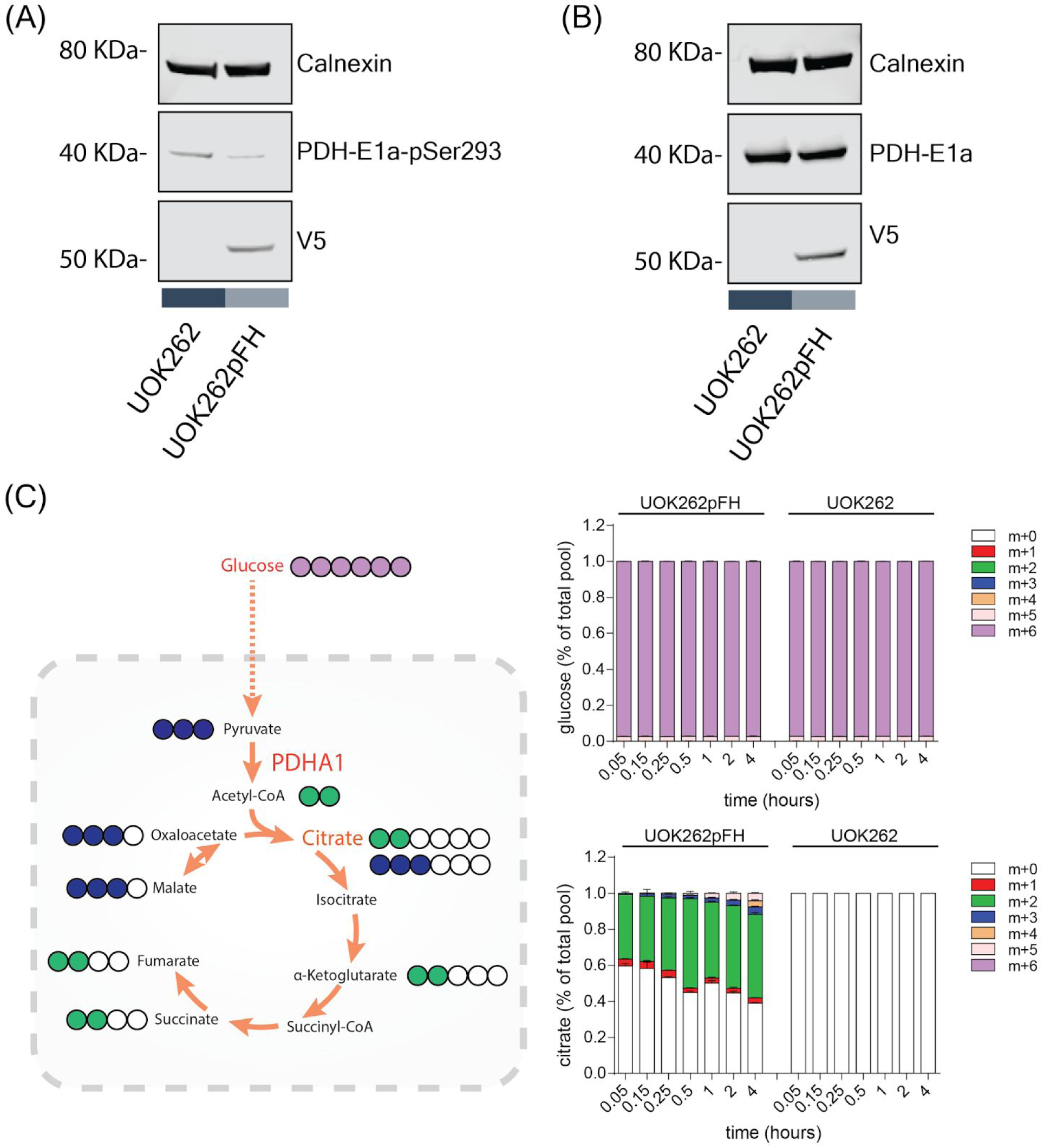
Experimental validation of PDHA1 phosphorylation regulation in FH-deficient cells. A) Western Blot of PDHA1 S293 using specific antibody. B) Western Blot of PDHA1 protein abundance. Calnexin was used as loading control and V5 to stain re-expressed V5-FH-wt in UOK262pFH. C) ^13^C-Glucose labelling experiment tracking the uptake of glucose into the mitochondria via PDHA1.

## Discussion

Metabolism deregulation is a hallmark of cancer (*2*,*25*), and it has become increasingly clear that it is a complex process involving post-transcriptional regulation, and understanding it is a challenging task that requires an integrative perspective. Here we acquired and analysed data characterizing the proteome, phosphoproteome, and metabolome of FH-deficient cell lines and found profound alterations that are impinged by restoring FH activity (Figure 1). We found strong down-regulation of the mitochondrial enzymes and significant phosphorylation changes in 23 phosphosites in metabolic enzymes. These findings suggest that phosphorylation regulates the activity of enzymes involved in central metabolic pathways, such as glycolysis and citric acid cycle, supporting the metabolic rewiring observed in FH-deficient cells.

Only a fraction of the phosphorylation-sites identified have a significant biological role (*26*) and as few as 20% are mapped to kinases or phosphatases (*20*, *27*). This observation emphasises the importance of assessing the functional impact of phosphorylation changes. Yet, it is challenging to predict how phosphorylation affects the activity of metabolic enzymes. In this work, we provided evidence that combining phosphoproteomics data with measurements of fluxes of metabolic enzymes enables an accurate prediction of how enzyme activity can be regulated by post-translational modifications. To estimate intracellular metabolic fluxes, we took advantage of genome-scale metabolic models constrained with quantitative experimental data of CORE rates of metabolites. Accurate quantification of the metabolic rates was important to rule out possible confounding effects, such as cell number and cell size, and to robustly estimate the metabolic differences between the two cell lines. The *in silico* estimated metabolic fluxes recapitulated several biological phenotypes of the UOK262 cell lines, for example, increased glycolytic flux, decreased mitochondrial respiratory function and increased lactate secretion (*10*). In general, the metabolic pathways fluxes were not associated with protein abundance, in agreement with results in microorganisms that showed that transcriptional profile is a poor predictor of metabolism (*18*). These results emphasise the importance to take in consideration other sources of regulation, in particular post-translational modifications such as phosphorylation.

Using the metabolic fluxes as readouts of functional activity of the metabolic enzymes, we performed a systematic identification of putative regulatory phosphosites of metabolic enzymes by matching phosphorylation changes in the enzymes residues with the flux changes of the catalysed reactions. Phosphorylation changes displayed strong correlation with protein abundance, and we therefore focused on metabolic enzymes that did not have significant protein abundance changes. From our list of putative regulatory phosphosites we recapitulated three, PDHA1_S232, PGK1_S203 and RRM2_S20, that were previously reported in PhosphositePlus (*20*). In particular, we explored the potential regulatory implication of S232 in FH-deficient cells. We validated the prediction that phosphorylation changes are also visible in another regulatory phosphosite of PDHA1, S293, and that decreased phosphorylation is accompanied by an increased flux of glucose into the mitochondria. PDHA1 functional phosphosites are regulated by the kinases PDK and phosphatases PDPK (*21*–*23*). While dysregulation of these proteins can provide mechanistic insight into the regulation of PDHA1, further work would need to be carried out, such as perturbation dynamic phosphoproteomics experiments, to validate the regulatory role of PDK and PDPK and of other potential upstream kinases and signalling pathways involved in FH-deficient cells. Another potential regulatory phosphosite is S83 in glyceraldehyde-3-phosphate dehydrogenase (GAPDH) for which our measurements show a significant decrease in phosphorylation and increase in metabolic flux. While further experimental evidence is required to confirm these findings, this hypothesis can potentially provide insights into the inactivation of GAPDH to re-route glycolysis flux into pentose phosphate pathway in response to potential oxidative stress (*28*) a phenomenon that has been reported in FH deficient cells (*29*).

In summary, this work provides for the first time a genome-scale study of the regulatory implications of post-translational modifications in the metabolism of FH-deficient cancer cell lines. Specifically, we exemplify the utility of our approach to identify in a systematic manner potential regulatory phosphorylation residues in metabolic enzymes, which can help to shed light into the complex regulation of cancer metabolism. This approach is also generally applicable to the study of other types of post-translational modifications that fall on metabolic enzymes, for example acetylation and succination, a modification caused by increased fumarate in FH-deficient cells (*30*). Pairing recent studies that have characterised the phosphoproteome of several hundreds of tumour samples (*31*, *32*) with metabolomics measurements will potentiate the discovery of novel therapies that exploit ubiquitous features of cancer metabolism.

## Methods

### UOK262 and UOK262pFH cells growth and sample preparation

Human FH-mutant UOK262 and FH-reconstituted UOK262pFH cells were obtained as previously described (*10*, *11*). All cells were grown in DMEM (Gibco 41966-029) supplemented with 10% heat inactivated FBS (Gibco 10270-106).

### Proteomics and phosphoproteomics mass-spectrometry experiment

Proteomics experiments were performed using mass spectrometry as reported (*33*, *34*). Urea lysis buffer was used to lyse the cells (8 M urea, 10 mM Na3VO4, 100 mM β-glycerol phosphate and 25 mM Na2H2P2O7 and supplemented with phosphatase inhibitors (Sigma)). Proteins were reduced and alkylated by sequential addition of 1 mM DTT and 5 mM iodoacetamide, followed by overnight incubation with immobilized trypsin to digest proteins into peptides. Using OASIS HLB columns (Waters) in a vacuum manifold peptides were desalted by solid phase extraction (SPE) following the manufacturer’s guidelines apart that the elution buffer contained 1 M glycolic acid. TiO_2_ enrichment beads (GL Sciences) was used for the phosphoproteomics analysis, similarly to that described (*34*, *35*) with minor modifications. Samples were analysed with LC-MS/MS using a LTQ-Orbitrap mass-spectrometer. Mascot was used to identify peptides against SwissProt human protein database, and Pescal used for quantification as previously described (*34*).

Nanoflow LC–MS/MS in an LTQ-orbitrap, as described in (*33*, *34*), was used to analyse dried peptide extracts dissolved in 0.1% TFA. 2% to 35% gradient elution from buffer B in 90 min with buffer A being used to balance the mobile phase (buffer A was 0.1 % formic acid in water and B was 0.1 % formic acid in acetonitrile). Mascot Distiller (version 1.2) was used to convert raw MS files (multistage acquisition) into Mascot Generic Format. SwissProt (version 2013.03) was used to search the MS with mass window of 10 ppm and 600 mmu for parent and fragment mass to charge values. Only human entries were considered using Mascot search engine (version 2.38). Searches for variable modifications were constituted by oxidation of methionine, pyro-glu (N-term) and phosphorylation of serine, threonine and tyrosine. False discovery rate of less than 1%, calculated by comparing against decoy databases, was considered. Peak areas of the first three isotopes of each peptide ion extracted chromatographs (mass 7 ppm and 1.5 min retention time window) was used for quantification. Retention shifts were taken in consideration by re-calculating for each peptide in each LC–MS/MS run individually using linear regression based on common ions across runs. The mass spectrometry proteomics data have been deposited to the ProteomeXchange Consortium via the PRIDE (*36*) partner repository with the dataset identifier PXD006693.

Raw intensities were log2 transformed followed by a scaling in each sample to account for potential pipetting differences. Only proteotypic peptides and peptides measured consistently across half of the replicates were considered. Differential protein abundance and phosphorylation analysis was performed using T-test between the UOK262 and UOK262pFH samples. P-values were controlled for false discovery rate using Benjamini–Hochberg FDR.

### Consumption and release quantification of metabolites

UOK262 and UOK262pFH (1.5x10^5^) were plated onto 6-well plates and allow to grow for 16h. 200 μl of cell culture media were collected from each well immediately after (t=0) and after additional 24 hours of incubation. Cells from the same plate were used for counting and then lysed in RIPA buffer for measurement of protein content. The collected media were centrifuged at 4°C for 10 min at max speed and 50 μl of the supernatant extracted in 750 μl of cold metabolites extraction buffer (MEB) as previously described (*10*). The solution was centrifuged at 4°C for 10 min at max speed and the supernatant was transferred onto LC-MS vials for metabolomics analyses.

LC-MS analysis of sample extracts was performed on a Q Exactive mass spectrometer coupled to Dionex UltiMate 3000 Rapid Separation LC system (Thermo). The liquid chromatography system was fitted with a SeQuant ZIC-pHILIC (150mm × 2.1mm, 5um) with guard column (20mm × 2.1mm, 5um) from Merck (Darmstadt, Germany). The mobile phase was composed of 20mM ammonium carbonate and 0.1% ammonium hydroxide in water (solvent A), and acetonitrile (solvent B). The flow rate was set at 180 uL x min-1 with the following gradient: 0 min 70% B, 1 min 70% B, 16 min 38% B, 16.5 min 70% B, hold at 70% B for 8.5 min. The mass spectrometer was operated in full MS and polarity switching mode. Medium from five independent cell cultures were analysed for each condition and samples were randomised in order to avoid bias in sample analyses due to machine drift. The acquired spectra were analysed using XCalibur Qual Browser and XCalibur Quan Browser software (Thermo Scientific) by referencing to an internal library of compounds.

Absolute quantification of metabolites in the cell culture medium was performed by interpolation of the corresponding standard curves obtained from commercially available compounds running with the same batch of samples. For each spent medium sample and each metabolite, the measured concentration spent was converted to consumption/release (CORE) data (molar amounts per dry weight per unit time) adapting the approach described in (*37*).

### Metabolic extracts after glucose and glutamine labelling

UOK262 and UOK262pFH (1.5x10^5^) were plated onto 6-well plates and grown overnight. The day after, medium was replaced with fresh one containing ^13^C_6_ glucose or ^13^C_5_ glutamine (Cambridge Isotope Laboratories). The following day, cells were counted using Countess (Thermo Fisher Scientific) and extracted as described before (*10*) at the indicated time points.

### Metabolic modelling using Recon 2.2

For this analysis the human metabolic reconstruction Recon 2.2 was used (*14*). The commonly used ATP maintenance (ATPM) reaction is added to the model, M_h2o_c + M_ atp_c → M_adp_c + M_pi_c + M_h_ c. Two context-specific models are generated: one for UOK262 cells and another for UOK262pFH. UOK262 model contains the specific measured growth rate, 0.01973, and hard coded deletion of FH by setting the upper and lower bounds of reactions FUM and FUMm to zero. UOK262pFH model is only constrained with the measured growth rate, 0.02940, since FH expression and activity is restored in these cell lines the catalysed reactions are not constrained. Uptake rates for all metabolites are constrained to zero, apart of those metabolites that present in cell culture medium, DMEM (Gibco 41966-029). For metabolites in the medium that were not measured or do not display significant CORE differences the exchange reactions lower bound were set similarly to the following publication (*38*). CORE measurements are used to constrain the metabolic models of UOK262 and UOK262pFH independently using a linear implementation of minimization of metabolic adjustment (MOMA) (*39*). The models are then simplified using flux variability analysis (FVA), thereby removing any reaction that is not capable of carrying flux. Models are simulated using pFBA (*15*) by maximising ATP production (ATPM reaction).

### Cell Lysates and western blot

6x10^5^ UOK262 and UOK262pFH cells were plated onto 6-cm dishes. After 24h, cells were washed twice in PBS on ice and then lysed using RIPA buffer. Protein content was quantified using Pierce BCA protein Assay (Thermo Fisher Scientific) following manufacturer’s protocol. 100 μg of proteins was heated at 70°C for 10 min in Bolt Loading Buffer 1x+4% β-mercapto-ethanol and then loaded onto a 4-12% Bolt Bis-Tris gel (Thermo Fisher Scientific). Gels were run at 165V using Bolt MES1x buffer for 40 minutes. Dry transfer of the proteins onto nitrocellulose membrane was obtained using IBLOT2 (Thermo Fisher Scientific). Membrane was then blocked for 1h at room temperature using in BSA or milk 5% in TBS 1X supplemented with Tween20 0.01% (TBST). Primary antibody for Calnexin (Abcam), V5 (Thermo Fisher Scientific), PDH-E1a (Thermo Fisher Scientific) and PDH-E1a-pSer293 (Millipore) were incubated overnight at 4°C. The day after, the membrane was washed in TBST and then incubated with secondary antibodies for 1h at room temperature (LiCOR, conjugated with 680 or 800 nm fluorophores). After washes in TBST, images were taken using Image Studio Lite software.

### Code dependencies and availability

All the computational analysis were performed in Python version 2.7.10 and are available under GNU General Public License V3 as GitHub projects in the following url https://github.com/saezlab/hlrcc. Metabolic modelling and SBML import of Recon 2.2 was performed using python module Framed version 0.3.2 (*40*). Plotting was done using Python modules Matplotlib version 1.4.3 (*41*) and Seaborn version 0.7.0 (*42*). Generalised linear models were built using Python modules Sklearn version 0.17.1 (*43*) and Statsmodels version 0.6.1 (*44*). Python modules Scipy version 0.17.1 (*45*) and Numpy version 1.11.1 (*46*) were used to perform efficient numerical calculations and statistical analysis. Biological data analysis and structuring was carried out using Python module Pandas version 0.18.1 (*47*).

## Author Contributions

C.F. and J.S.R. conceptualised the project. E.G. carried the computational analysis. M.S., A.S.H.C. and T.I.J. performed the experimental work and validations. D.M. supervised the metabolic modelling. E.G. wrote the paper, with input from C.F. and J.S.R.

## Acknowledgments

We gratefully acknowledge helpful comments from Pedro Beltrao and David Ochoa.

## Supplementary figures

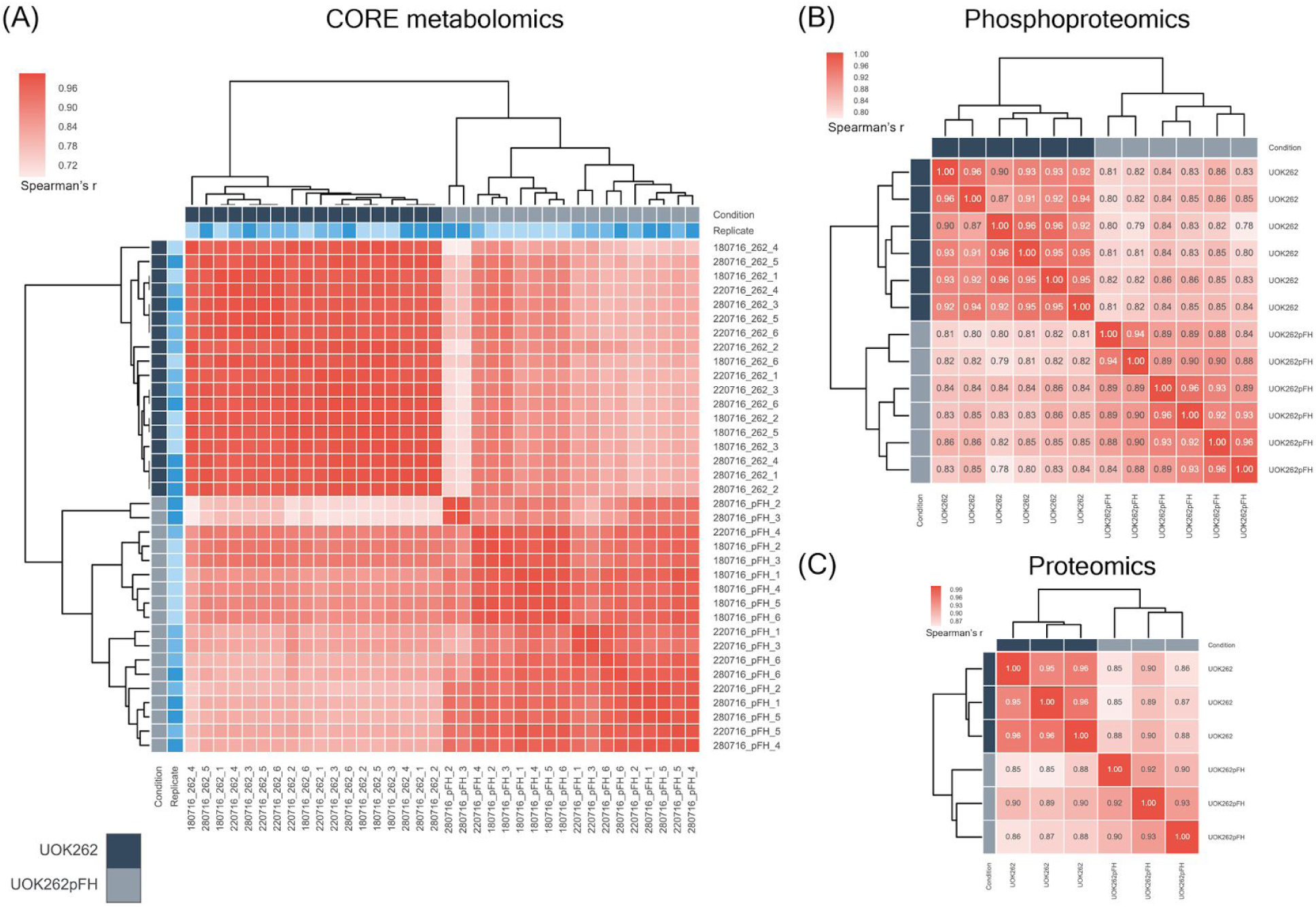
Supplementary Figure 1. Unsupervised clustering of the experimental replicates. A) Clustering analysis of the three different biological and technical replicates of the CORE metabolomics experiments. Light blue colour denotes the different biological replicates. B) Clustering of the three biological replicates, where each has a technical replicate, of the phosphoproteomics experiments. C) Clustering analysis of the three biological replicates of the proteomics measurements.

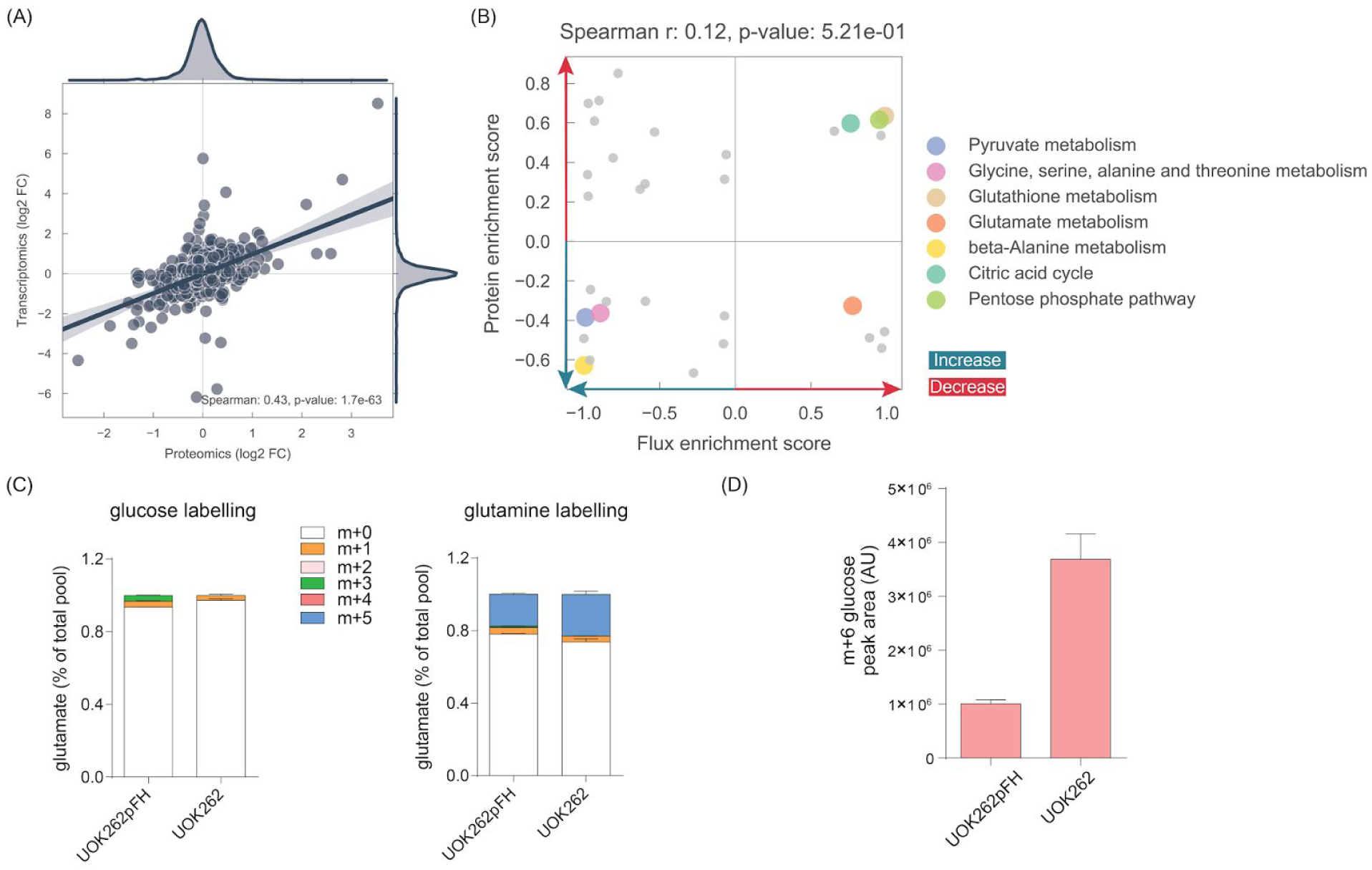
Supplementary Figure 2. Proteomics correlation analysis. A) Correlation between proteomics and transcriptomics measurements. Log2 fold changes between UOK262 and UOK262pFH were used. B) Scatter plot with the GSEA enrichment score obtained for each metabolic pathway in the metabolic model for which it was possible to overlap at least 5 measured proteins or *in silico* estimated fluxes. Representative pathways were highlighted. C) Glucose uptake in UOK262pFH and UOK262. D) Glutamate labelling from glucose or glutamine secreted in the medium of UOK262 and UOK262pFH. The time point used for glucose labelling is 4h while for glutamine is 3h.

### Supplementary Tables

**Supplementary Table 1.** Differential analysis of proteomics, phosphoproteomics and CORE metabolomics.

**Supplementary Table 2.** List of *in silico* estimated fluxes for UOK262 and UOK262pFH cells.

**Supplementary Table 3.** GSEA enrichment analysis output performed in both proteomics and fluxomics data-sets.

**Supplementary Table 4.** List of putative regulatory phosphorylation-sites located in metabolic enzymes.

## References

1. M. R. Stratton, P. J. Campbell, P. A. Futreal, The cancer genome. Nature. 458, 719–724 (2009).

2. D. Hanahan, R. A. Weinberg, Hallmarks of cancer: the next generation. Cell. 144, 646–674 (2011).

3. J. Hu et al., Heterogeneity of tumor-induced gene expression changes in the human metabolic network. Nat. Biotechnol. 31, 522–529 (2013).

4. E. Gaude, C. Frezza, Tissue-specific and convergent metabolic transformation of cancer correlates with metastatic potential and patient survival. Nat. Commun. 7, 13041 (2016).

5. A. P. Oliveira et al., Dynamic phosphoproteomics reveals TORC1-dependent regulation of yeast nucleotide and amino acid biosynthesis. Sci. Signal. 8, rs4 (2015).

6. Z. Raguz Nakic, G. Seisenbacher, F. Posas, U. Sauer, Untargeted metabolomics unravels functionalities of phosphorylation sites in Saccharomyces cerevisiae. BMC Syst. Biol. 10, 104 (2016).

7. E. Gonçalves et al., Systematic Analysis of Transcriptional and Post-transcriptional Regulation of Metabolism in Yeast. PLoS Comput. Biol. 13, e1005297 (2017).

8. I. P. M. Tomlinson et al., Germline mutations in FH predispose to dominantly inherited uterine fibroids, skin leiomyomata and papillary renal cell cancer. Nat. Genet. 30, 406–410 (2002).

9. J. S. Isaacs et al., HIF overexpression correlates with biallelic loss of fumarate hydratase in renal cancer: novel role of fumarate in regulation of HIF stability. Cancer Cell. 8, 143–153 (2005).

10. M. Sciacovelli et al., Fumarate is an epigenetic modifier that elicits epithelial-to-mesenchymal transition. Nature. 537, 544–547 (2016).

11. C. Frezza et al., Haem oxygenase is synthetically lethal with the tumour suppressor fumarate hydratase. Nature. 477, 225–228 (2011).

12. N. C. Duarte et al., Global reconstruction of the human metabolic network based on genomic and bibliomic data. Proc. Natl. Acad. Sci. U. S. A. 104, 1777–1782 (2007).

13. I. Thiele et al., A community-driven global reconstruction of human metabolism. Nat. Biotechnol. 31, 419–425 (2013).

14. N. Swainston et al., Recon 2.2: from reconstruction to model of human metabolism. Metabolomics. 12, 1–7 (2016).

15. N. E. Lewis et al., Omic data from evolved E. coli are consistent with computed optimal growth from genome-scale models. Mol. Syst. Biol. 6, 390 (2010).

16. A. Subramanian et al., Gene set enrichment analysis: a knowledge-based approach for interpreting genome-wide expression profiles. Proc. Natl. Acad. Sci. U. S. A. 102, 15545–15550 (2005).

17. The Gene Ontology Consortium, Gene Ontology Consortium: going forward. Nucleic Acids Res. 43, D1049–D1056 (2015).

18. D. Machado, M. Herrgård, Systematic evaluation of methods for integration of transcriptomic data into constraint-based models of metabolism. PLoS Comput. Biol. 10, e1003580 (2014).

19. L. Zheng et al., Reversed argininosuccinate lyase activity in fumarate hydratase-deficient cancer cells. Cancer Metab. 1, 12 (2013).

20. P. V. Hornbeck et al., PhosphoSitePlus, 2014: mutations, PTMs and recalibrations. Nucleic Acids Res. 43, D512–20 (2015).

21. L. G. Korotchkina, M. S. Patel, Site specificity of four pyruvate dehydrogenase kinase isoenzymes toward the three phosphorylation sites of human pyruvate dehydrogenase. J. Biol. Chem. 276, 37223–37229 (2001).

22. M. Kato et al., Structural basis for inactivation of the human pyruvate dehydrogenase complex by phosphorylation: role of disordered phosphorylation loops. Structure. 16, 1849–1859 (2008).

23. F. Seifert et al., Phosphorylation of serine 264 impedes active site accessibility in the E1 component of the human pyruvate dehydrogenase multienzyme complex. Biochemistry. 46, 6277–6287 (2007).

24. Y. Yang et al., Metabolic reprogramming for producing energy and reducing power in fumarate hydratase null cells from hereditary leiomyomatosis renal cell carcinoma. PLoS One. 8, e72179 (2013).

25. D. Hanahan, R. A. Weinberg, The hallmarks of cancer. Cell. 100, 57–70 (2000).

26. P. Beltrao et al., Systematic functional prioritization of protein posttranslational modifications. Cell. 150, 413–425 (2012).

27. H. Dinkel et al., Phospho. ELM: a database of phosphorylation sites—update 2011. Nucleic Acids Res. 39, D261–D267 (2011).

28. C. M. Grant, Metabolic reconfiguration is a regulated response to oxidative stress. J. Biol. 7, 1 (2008).

29. C. Sourbier et al., Targeting ABL1-Mediated Oxidative Stress Adaptation in Fumarate Hydratase-Deficient Cancer. Cancer Cell. 26, 840–850 (2014).

30. M. Yang et al., The Succinated Proteome of FH-Mutant Tumours. Metabolites. 4, 640–654 (2014).

31. H. Zhang et al., Integrated Proteogenomic Characterization of Human High-Grade Serous Ovarian Cancer. Cell. 166, 755–765 (2016).

32. P. Mertins et al., Proteogenomics connects somatic mutations to signalling in breast cancer. Nature. 534, 55–62 (2016).

33. V. Rajeeve, I. Vendrell, E. Wilkes, N. Torbett, P. R. Cutillas, Cross-species proteomics reveals specific modulation of signaling in cancer and stromal cells by phosphoinositide 3-kinase (PI3K) inhibitors. Mol. Cell. Proteomics. 13, 1457–1470 (2014).

34. P. Casado et al., Kinase-substrate enrichment analysis provides insights into the heterogeneity of signaling pathway activation in leukemia cells. Sci. Signal. 6, rs6 (2013).

35. A. Montoya, L. Beltran, P. Casado, J.-C. Rodríguez-Prados, P. R. Cutillas, Characterization of a TiO2 enrichment method for label-free quantitative phosphoproteomics. Methods. 54, 370–378 (2011).

36. J. A. Vizcaíno et al., 2016 update of the PRIDE database and its related tools. Nucleic Acids Res. 44, D447–56 (2016).

37. M. Jain et al., Metabolite profiling identifies a key role for glycine in rapid cancer cell proliferation. Science. 336, 1040–1044 (2012).

38. K. Yizhak, O. Gabay, H. Cohen, E. Ruppin, Model-based identification of drug targets that revert disrupted metabolism and its application to ageing. Nat. Commun. 4, 2632–2632 (2013).

39. D. Segrè, D. Vitkup, G. M. Church, Analysis of optimality in natural and perturbed metabolic networks. Proc. Natl. Acad. Sci. U. S. A. 99, 15112–15117 (2002).

40. D. Machado, Framed (2017; https://zenodo.org/record/240430).

41. J. D. Hunter, Matplotlib: A 2D Graphics Environment. Computing in Science Engineering. 9, 90–95 (2007).

42. M. Waskom et al., seaborn : v0.5.0 (November 2014) (ZENODO, 2014; http://zenodo.org/record/12710).

43. F. Pedregosa et al., Scikit-learn: Machine Learning in Python. J. Mach. Learn. Res. 12, 2825–2830 (2011).

44. Statsmodels (2009), (available at http://statsmodels.sourceforge.net).

45. E. Jones, T. Oliphant, P. Peterson, Others, SciPy: Open source scientific tools for Python (2016), (available at http://www.scipy.org/).

46. S. van der Walt, S. C. Colbert, G. Varoquaux, The NumPy Array: A Structure for Efficient Numerical Computation. Computing in Science Engineering. 13, 22–30 (2011).

47. W. McKinney, Others, in Proceedings of the 9th Python in Science Conference (2010), vol. 445, pp. 51–56.

